# The relationship between microbial population size and disease in the *Arabidopsis thaliana* phyllosphere

**DOI:** 10.1101/828814

**Authors:** Talia L. Karasov, Manuela Neumann, Alejandra Duque-Jaramillo, Sonja Kersten, Ilja Bezrukov, Birgit Schröppel, Efthymia Symeonidi, Derek S. Lundberg, Julian Regalado, Gautam Shirsekar, Joy Bergelson, Detlef Weigel

## Abstract

A central goal in microbiome research is to learn what distinguishes a healthy from a dysbiotic microbial community. Shifts in diversity and taxonomic composition are important indicators of dysbiosis, but a full understanding also requires knowledge of absolute microbial population sizes. In addition to the number of microbial cells, information on taxonomic composition can provide important insight into microbiome function and disease state. Here we use shotgun metagenomics to simultaneously assess microbiome composition and microbial load in the phyllosphere of wild populations of the plant *Arabidopsis thalian*a. We find that wild plants vary substantially in the load of colonizing microbes, and that high loads are typically associated with the proliferation of single taxa, with only a few putatively pathogenic taxa achieving high abundances in the field. Our results suggest (i) that the inside of a plant leaf is on average sparsely colonized with an estimated two bacterial genomes per plant genome and an order of magnitude fewer eukaryotic microbial genomes, and (ii) that higher levels of microbial cells often indicate successful colonization by pathogens. Lastly, our results show that load is a significant explanatory variable for loss of estimated Shannon diversity in phyllosphere microbiomes, implying that reduced diversity may be a significant predictor of microbial dysbiosis in a plant leaf.

## Introduction

Host-associated microbes can be classified according to the effects they have on the host, with some benefitting the host [1], others hurting the host [2], and most having no apparent impact. For only a small subset of specific microbes do we know to which of these categories they belong. Individual testing of thousands of microbial taxa in physiologically relevant settings remains infeasible for most host-microbe systems. A key challenge in understanding microbiome function is therefore the development of methods that help us to infer indirectly which of the numerous taxa associated with a host also have a measurable impact on host health.

In both plants and animals, compositional microbiome analyses such as 16S rDNA amplicon sequencing have been integral to revealing the hundreds or even thousands of microbial species that co-occur within single tissues, and the responses of taxa to environmental differences [3–5]. While compositional analyses indicate changes in the number of microbial cells relative to each other, these data alone do not tell us whether a focal taxon changed in absolute abundance relative to the host. A proxy for what we term here the microbial load of a host is therefore the ratio of microbial to plant cells, and this may in turn be an important indicator of host health. Indeed, assessing the inability of a host to control pathogen proliferation is a common metric for assessing disease state [6], and the importance of evaluating microbial load is increasingly acknowledged in microbiome studies. For example, a recent study demonstrated that inferring correlations between human disease and microbial taxa is contingent on microbial load in the gut [7]. In probing the relationship between microbiome composition, microbial load, and Crohn’s disease, the authors found that only when microbial load was taken into account was it possible to detect associations between specific taxa and disease state. There are also examples of low biomass microbes exerting a significant effect on host health [8]. Therefore, at least in humans, and likely in other mammals as well, information on microbial load is apparently informative for identifying novel disease agents and dysbiotic microbial communities.

In plants, microbial load has been shown to differ substantially between individuals of the same and different species. Maignien and colleagues [9] found bacterial load in naturally colonized *Arabidopsis thaliana* populations to vary by an order of magnitude even within a single population. Karasov and colleagues [2] similarly found that wild *A. thaliana* in the same local metapopulation differed by an order of magnitude in their estimated microbial loads, and discerned a possible association between microbial load and putative infection. The major driver of differences in microbial load was a pathogenic *Pseudomonas* taxon, which indicated that this taxon may have reduced the ability of the host to control bacterial growth in its leaves. These studies raise the possibility that microbial load could be used to identify–without prior information–taxa that are novel disease agents.

At present, we lack a baseline for how many microbial cells on average colonize the leaf of a wild plant. Only once this baseline is established (or at least approximated) can we determine its relationship to host health. With this information we can, for example, identify exceptional plants with abnormally high or low loads, and analyze these individuals in detail to discover host, microbial and environmental factors that influence load.

In this study, we make use of data on the progression of infection in the laboratory to inform our understanding of microbiome colonization in wild plant populations. We begin with controlled infections of a bacterial and an oomycete pathogen, and monitor the microbial load and composition throughout disease progression. We then analyze the relationship between microbial load, composition, and diversity in the phyllosphere of wild *A. thaliana* populations in Germany and Sweden. The two geographic regions provide contrasts in both microbiome composition and load, with higher loads being due primarily to the proliferation of putatively pathogenic taxa of both bacteria and oomycetes. Using the controlled infections as standards, we find that in wild populations only a few taxa proliferate to a load higher than basal colonization in a laboratory-grown plant. Importantly, these high-load taxa encompass the most common pathogens of *A. thaliana*, such as *Pseudomonas* sp. [10] and *Hyaloperonospora arabidopsidis [11]*. Based on our observations, we propose that high microbial population size in the *A. thaliana* phyllosphere may be achieved nearly exclusively by pathogenic taxa. This prediction can now be tested by the systematic analysis of other *A. thaliana* populations.

## Results

### Microbial load correlates with disease under laboratory conditions

Molecular plant pathology, especially in *A. thaliana*, has a long history of assessing the pathogen colonization process through laboratory infections [12]. This progression is typically tracked by infecting plants with a low titer of pathogen, then following the growth of the pathogen via plating or quantification of microbial growth via microscopy [13]. To assess the level of colonization of a host by a known bacterial pathogen, one can culture the pathogen and count colonies. For eukaryotic filamentous pathogens such as fungi and oomycetes, one can quantify hyphae and spores by microscopy. Alternatives for determining the overall load of microbes are quantitative polymerase chain reaction (qPCR) [7], spike-ins of known taxa to extracted DNA [14], and shotgun metagenomics [15, 16]. While every method of estimating microbial load has its limitations [17, 18], among approaches that can be easily scaled to a large number of samples, shotgun metagenomics has several advantages: it avoids many well-established biases in amplicon sequencing [19, 20], and it has the potential to simultaneously provide information on community composition and microbial population sizes. We sought to create from laboratory infections the equivalent of metagenomics standard curves that had the potential to inform subsequent observations of wild plants. To this end, we performed controlled infections with a model prokaryotic pathogen (*P. syringae*) under standard laboratory conditions, followed by simultaneously measuring disease progression over time with traditional plant pathology metrics (colony forming units), qPCR measures of microbial abundance, and metagenomic load quantification by whole-genome shotgun sequencing. We previously demonstrated that one can obtain quantitative measures of microbial colonization in wild plants by performing shotgun metagenomic sequencing of whole plant tissue, followed by determining the ratio of reads assigned as having either a microbial or plant origin [2]. We have further validated our approach by showing that compositional inferences agree well between this method and more conventional amplicon sequencing methods [16].

For the *P. syringae* analysis, we performed controlled infections in the laboratory of the *A. thaliana* reference accession Col-0 with two *P. syringae* strains: Pst DC3000:avrB, which is recognized by the host immune system of this accession [21], and Pst DC3000:EV, which is not [22]. The formal plant pathology terms for these are incompatible and compatible interactions [23, 24]. Both *P. syringae* strains carry the full effector complement required for successful colonization of *A. thaliana*.

With these *P. syringae* strains, we performed infections of Col-0 with a Pst DC3000 [25] inoculum often found in the plant pathology literature (OD_600_=0.0002 or approximately 105 colony forming units [cfu] per mL) [26], and collected samples over six days (Figure 1). We syringe-inoculated plants and took daily samples of 5-6 replicates. With each sampling, we performed colony counting on extracts from leaf hole punches to directly measure live bacterial content, we performed qPCR to quantify the ratio of plant DNA to the 16S rDNA of colonizing bacteria, and we performed shotgun metagenomics and taxonomic assignment to assess the ratio of plant metagenomic reads to Pseudomonadaceae metagenomic reads.

**Figure 1:**
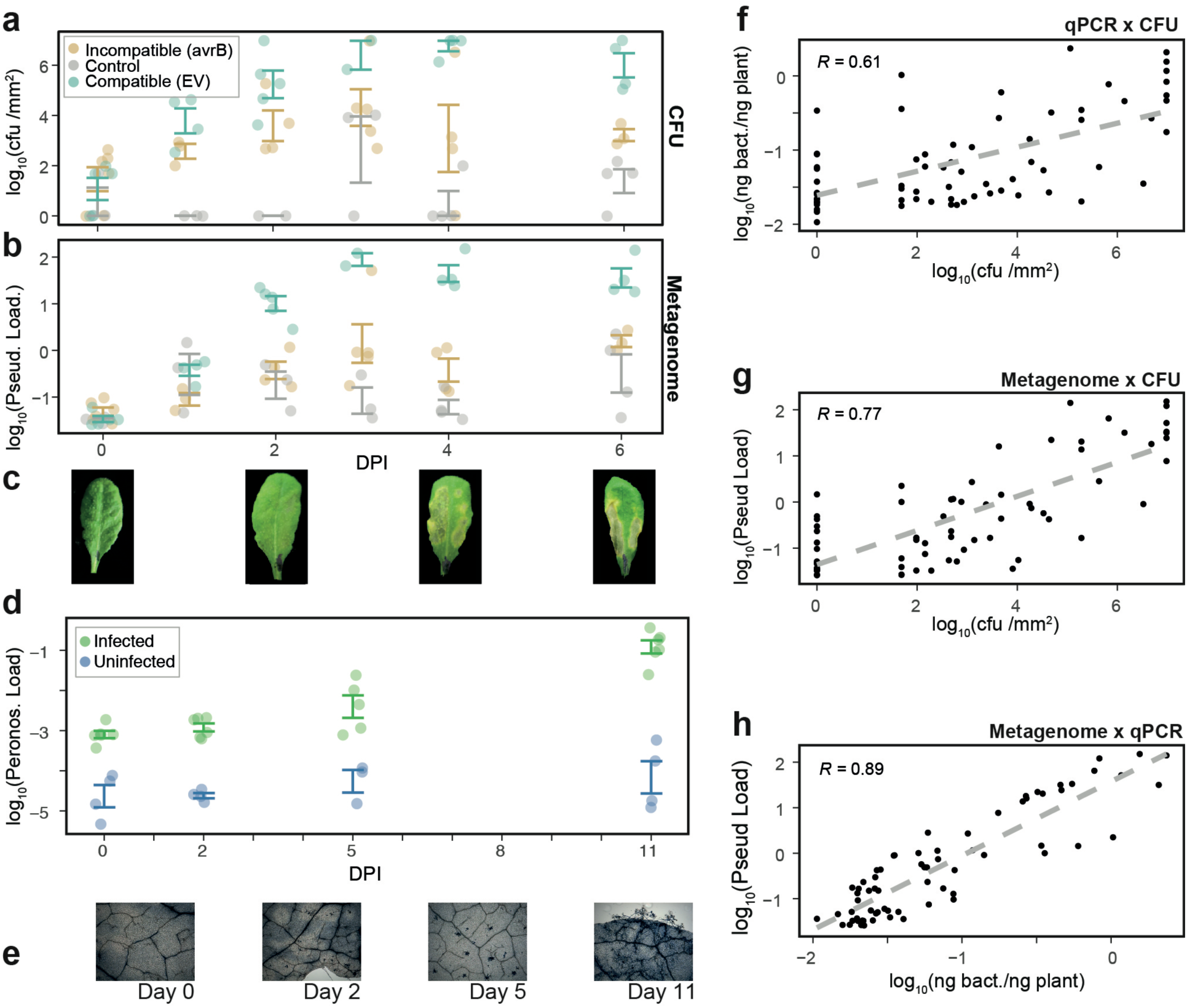
Comparison of different metrics of *P. syringae* and *HpA* growth *in planta*. (a) Time-series of *P. syringae* growth (measured in colony forming units [CFU] per mm^2^).Time was measured as days post infection (DPI) (b) Time-series of metagenomic load of Pseudomonadaceae (measured as ratio of coverage of Pseudomonadaceae genome to coverage of *A. thaliana* genome). (c) Representative images of infection progression with Pst DC3000 in Col-0. (d) Time-series of Peronosporaceae load (measured as ratio of coverage of Peronosporaceae genome to coverage of *A. thaliana* genome) throughout infection progression. Data represented as mean values +/- standard error. (e) Representative images of *HpA*-infected samples after Trypan blue staining of dead tissue and *HpA* structures. (f) Scatterplot of relationship between CFU (x-axis) and qPCR measurement (y-axis), (g) CFU (*x*-axis) and metagenomic measurement (*y*-axis), and (h) the relationship between qPCR (*x*-axis) and metagenomic measurement (*y*-axis).The figures show the line of best fit in linear regression with log-transformed data. Related to Figure S1.

Qualitatively, the three parallel measures of Pst DC3000 growth over time were remarkably similar, with an increase in bacterial abundance for the first three days after infection, followed by a plateau in microbial propagation (Figure 1a). Comparisons of the log-transformed measurements of the data confirmed that qPCR and metagenomic load measurements were significantly correlated (r = 0.88, p < 2×10^−16^) (Figure S1), and that the correlation coefficient between metagenomic load measurements and live colony counting (r = 0.77, p < 2×10^−16^) was higher than the correlation coefficient between qPCR measurements and colony counting (r = 0.61, p < 2×10^−16^). This may be because the qPCR amplification method we used did not distinguish between the 16S rDNA amplicons of Pseudomonadaceae and those of other bacterial families, whereas the metagenomic classification did. We also observed the presence of Pseudomonadaceae in the control infections for all measurement methods. This is most likely due to the abundance of Pseudomonads in most soils and non-sterile lab-grown plants [3]. Regardless, the metagenome-based metric of load quantification and qPCR method were consistent with one another.

Eukaryotic microbes are also frequent pathogens of *A. thaliana* and among the most common ones is *HpA* [27]. An obligate biotroph, *HpA* spreads between plants as spores [11]. Under humid conditions, the spores will develop hyphae, which penetrate the plant epidermis and subsequently the mesophyll (Figure 1), eventually form fruiting bodies. These fruiting bodies then sporulate, releasing the spores to infect other plant tissues. Assessing the abundance of *HpA* via nucleic acid-based methods is complicated by the fact that its hyphae are multinucleated [28]. Consequently, one “cell” of *HpA* encapsulates more than one nuclear genome.

For the *HpA* analysis, we took advantage of an *HpA* strain that we recently collected and an *A. thaliana* genotype related to those from which this *HpA* strain was isolated. We drip-inoculated the *A. thaliana* genotype H2081 with the *HpA* isolate 14OHML004 (both from the Midwestern USA), then took samples of plants over an eleven-day time window. Microbial load as determined by shotgun metagenomics was used as parallel metric to assess the growth of *HpA* and was compared with the qualitative images of hyphal growth visualized using Trypan blue staining [29].

Sporulation commenced around day 5 of infection, as seen both in the Trypan blue staining and in the substantial increase in microbial load by day 11. Until sporulation was induced on day 5, the microbial load of *HpA* was low (approximately three *HpA* genomes per 1,000 *A. thaliana* genomes). After induction, the load increased to an average of seven *HpA* genomes per 100 *A. thaliana* genomes.

Overall, our results obtained by whole-genome shotgun sequencing were consistent with many previous reports on laboratory infections with the model pathogens Pst DC3000 and *HpA*. In addition, by obtaining more quantitative measures of absolute abundance, we could conclude that Pst DC3000, a prokaryotic microbe, can achieve microbial loads–on a chromosome-per-chromosome level–that are more than an order of magnitude higher than *HpA*, a eukaryotic microbe, can.

### *Pseudomonas* load is negatively associated with plant growth

Previous studies identified differences in load across plant populations. While it was intriguing to observe overall differences in microbial loads across plants, we also wanted to know whether increased microbial load, particularly of a known *A. thaliana* colonizer, is associated with decreased plant growth. Previous studies testing gradients of initial infection doses with a *P. syringae* pathogen demonstrated that higher initial infection titers led to more prominent symptoms and reduced plant fitness [30, 31]. To determine whether this relationship holds for other combinations of *Pseudomonas* and *A. thaliana* genotypes, we replicated the infection experiments described in ref. [2] with pairs of one of two *A. thaliana* genotypes and one of three *Pseudomonas* strains and simultaneously measured both plant and microbial growth (Figure S2). Importantly, we tested this relationship with strains of *Pseudomonas* isolated from *A. thaliana* and known to colonize to high titer in wild *A. thaliana* populations.

In these assays, we tagged the focal *Pseudomonas* strains with luciferase to track the growth of the bacteria, and detection of plant-associated green pixels in images of rosettes to track the growth of the plant. Across all *A. thaliana* and *Pseudomonas* genotypes, proliferation of *Pseudomonas* was negatively associated with the size of *A. thaliana* host plants, and significantly so for 4/6 genotype x genotype combinations (Wilcoxon rank-sum test of each plant-microbe interaction, p < 0.05 after Benjamin-Hochberg correction [32]). These results suggest that for the *Pseudomonas* strains we tested here higher bacterial loads are associated with reduced plant health, and likely reduced plant fitness. While this relationship could differ with other Pseudomonads, we note that the strains tested here belong to the most abundant taxonomic group (by both absolute number and relative abundance) of *Pseudomonas* in the *A. thaliana* phyllosphere in Southwestern Germany [2].

### Microbial load in nature differs within and between regions, but high levels of microbial colonization are rare

After having obtained data on the correlation between conventional measures of infection in the lab and microbial load as determined by whole-genome shotgun sequencing, we proceeded to apply our methodology to the endophytic compartments of the phyllosphere of wild *A. thaliana* plants.

To ensure that our results do not simply reflect the idiosyncrasies of a single geographic region, we targeted two regions with contrasting climates and environments: Southwestern Germany and Sweden (Figure S3, Table S1). Within each region we surveyed several populations. With this information our aim was to determine (i) which microbial families contribute the most to microbial load, (ii) whether the same families within a region are reproducibly associated with high load, (iii) whether there is a consistent relationship between load and microbial diversity, and (iv) whether loads observed in nature are similar to loads obtained after compatible or incompatible laboratory infections.

To maximize reproducibility across samples, all samples were processed in the same manner. A strength of our metagenomic comparison method is that we compare the ratios of abundances across samples, an approach that can bypass several known biases in analyzing shifts in microbiome composition [18]. Nonetheless, we recognize that a metagenomic assessment of microbial DNA is merely a proxy for microbial load, and not a direct measurement.

Our previous work demonstrated that plants within the same population differ in microbial load [2] and that the presence of a specific taxonomic group from the genus *Pseudomonas* is correlated with high loads in Southwestern Germany. We analyzed these previously published data from four populations in Southwestern Germany [2] in more detail. Using the same washing methods as before, we generated in addition new shotgun metagenomic data from washed leaves of *A. thaliana* plants collected from five natural populations in Sweden that differed from but were in proximity to *A. thaliana* populations studied and described elsewhere [33]. We mapped the shotgun metagenomic data to the *A. thaliana* Col-0 reference genome [34], and classified unmapped reads using the k-mer metagenomic mapping tool *centrifuge* [35]. As shown before, the coverage estimates are robust and largely insensitive to the *A. thaliana* reference genome used [16]. We then estimated the ratio of average coverage over microbial genomes to plant nuclear genomes.

We estimated the average total load of a plant, per 100 plant genomes, to be 250 (s.d. = 312) bacterial genomes, 1.14 (s.d. = 1.34) fungal genomes, and 3.3 (s.d. = 1.63) oomycete genomes (Figure 2-3).

**Figure 2:**
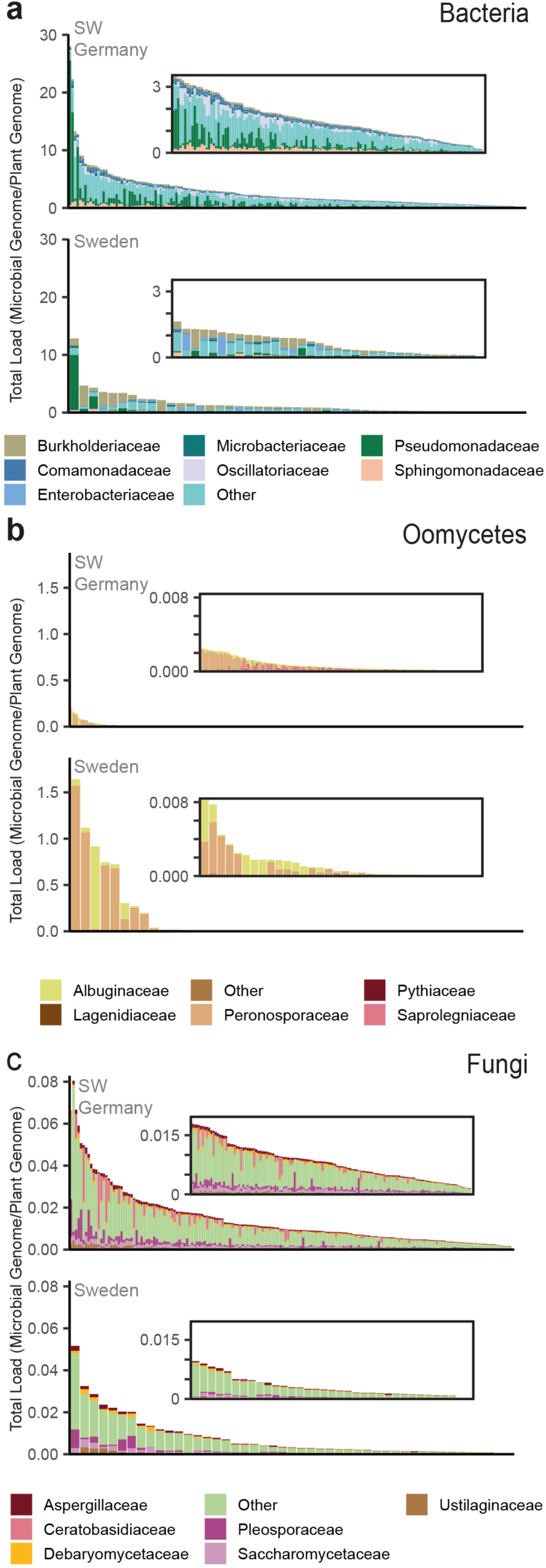
The microbial load and composition of *A. thaliana* populations in Germany and Sweden. (a)-(c) Bacterial, oomycete and fungal load and composition classified at the family taxonomic level for plants collected in Southwestern Germany and Sweden. Each figure shows the full distribution across all plants with an inset subplot.

**Figure 3:**
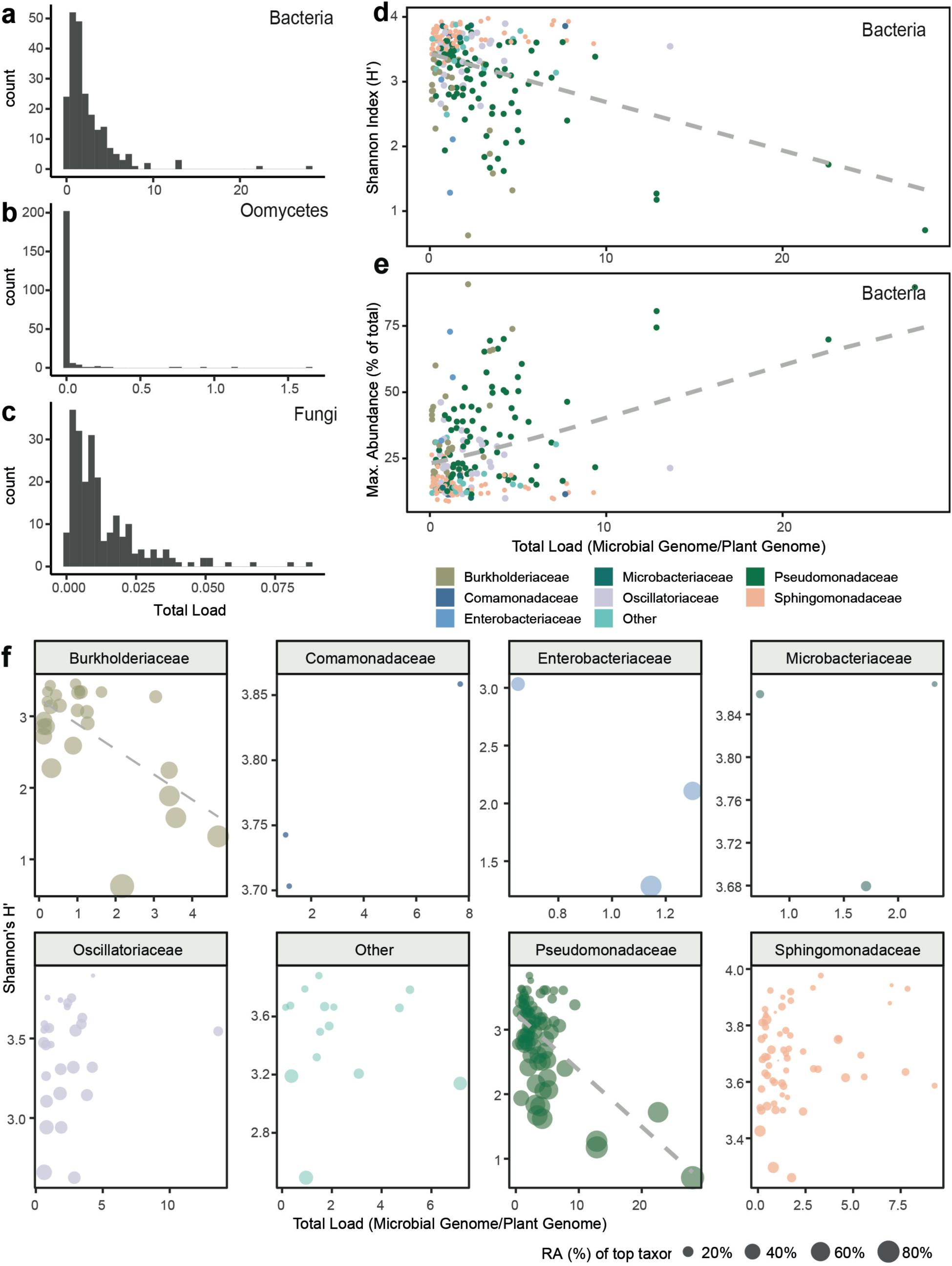
Microbial load is a significant predictor of microbiome diversity. Histogram of total load per plant of (a) bacteria (b) oomycetes and (c) fungi. (d) Total bacterial load is significantly negatively correlated with the Shannon’s H’. The dashed line is a line of best fit in linear regression for Shannon Diversity. Points are colored according to the most abundant taxon in a given sample. (e) Total bacterial load is positively associated with an increase in the abundance of a single family. The dashed line is a line of predicted values in beta regression with logit link (Ferrari and Cribari-Ne-to 2004). (f) A magnified illustration of (e) with each panel showing only those samples in which the focal taxon was the most abundant. The size of each point represents the percentage of the total load of a sample due to the focal taxon. Dashed lines are shown for the two families for which the coefficient estimate for Total Load in the linear regression of Shannon’s H’∼Total Load was significant (F-test, p < 0.05).

The samples from Germany consisted of 176 plants representing four populations over two seasons [2]. The samples from Sweden consisted of 46 plants collected from five locations in a single season (Figure S3). Differential abundance comparisons on variance-stabilized relative abundance data [36, 37] and ordination easily distinguished plants from Germany and Sweden based on their total microbial load and microbiome composition (Figure 2, S4-5). The bacterial, fungal, and oomycete families in Germany with the highest average loads were Pseudomonadaceae, Ceratobasidiaceae, and Peronosporaceae, while in Sweden they were Burkholderiaceae, Pleosporaceae, and Peronosporaceae. Compared to Swedish plants, German plants had on average higher bacterial load (Wilcoxon rank-sum test p = 8×10^−7^), and lower fungal load (Wilcoxon rank-sum test p = 0.006), but were not significantly different in oomycete load (Wilcoxon rank-sum test p = 0.251).

Considering only bacterial families, the metagenomes from the two regions were compositionally distinct (permutational analysis of variance [38] mean partial r^2^ = 0.27, p = 0.001). Indeed, 33/379 bacterial families differed in abundance by more than two-fold between the two regions (adjusted p-value [39] < 0.05, Wald-test), with twenty-three of those families more abundant in Germany than in Sweden. Compositional distinctness was true for populations in each region (mean partial r^2^ = 0.18, p = 0.001). Load was also a significant explanatory variable for differences in microbiomes across regions (PERMANOVA, mean partial r^2^ = 0.06, p = 0.008). Because the German and Swedish plants had been collected at different times and processed separately, we could not rule out that some of the observed differences between regions also reflected differences in sample processing. In light of this possibility, we focused on the population for which we had the most plants sampled (Eyach in Germany, n = 86) and tested within this population the relationship between load and the Bray-Curtis measure of community dissimilarity. Within the Eyach population alone, differences in load were also significantly associated with differences in composition (permutational analysis of variance r^2^ = 0.19, p = 0.001).

### High microbial load is associated with proliferation of single taxa and reduced taxonomic diversity

Our previous work indicated that high microbial load in German populations was correlated with the presence of strains representing a single pathogenic taxon of *Pseudomonas* [2]. To test whether taxa other than *Pseudomonas* could be responsible for high load, we calculated the relationship between the bacterial load of a plant leaf and the maximum relative abundance of a family observed per sample (Figure 3). There was a positive relationship between these two variables for bacteria (Beta regression Pseudo-R = 0.36, p < 4×10^−8^) and the results were robust to the exclusion of plants with the highest loads (load > 10).

Our results revealed that in the most heavily colonized plants, those with highest microbial load, a single taxonomic family of oomycetes or bacteria explained the majority of the load increase relative to less heavily colonized plants. A direct consequence of this relationship was a reduction in the estimated Shannon’s Diversity Index in the leaves of plants with a high load (Figure 3). Importantly, the reduced bacterial Shannon diversity was not an artifact of reduced depth of sequencing of bacterial reads (Figure S6). The reduced diversity could be the result of at least two distinct processes: (1) The most abundant family increases in abundance without an influence on the surrounding microbial presence, or (2) the most abundant family suppresses the surrounding microbiota while proliferating. We did not see a negative correlation between the load of Pseudomonadaceae and the load of any other bacterial families (Figure S6), indicating that the change in observed diversity is not due to the effect on the surrounding microbes and on species richness, but instead to the proliferation of Pseudomonadaceae.

That bacterial and oomycete families associated with high load contained well-known pathogens, such as *P. syringae* and *Hyaloperonospora arabidopsidis* (*HpA*), further suggesting that high load could be an indicator of plant disease state. We note that when plants were sampled, we had done so without regard to the presence or absence of visual signs of disease. In the field it is often difficult to assess disease state of *A. thaliana* plants, as there are numerous causes for reduced plant size and chlorosis, including non-optimal temperatures [40] and drought [41, 42]. Consequently, for field-collected plants, unless we have *a priori* information on the presence of disease agents, we can rarely assess microbe-associated disease state and progression directly.

### Few microbial taxa in wild populations exceed loads observed in incompatible infections

In contrast to disease assessment in the field, in standard laboratory infections it is relatively straightforward to assess disease state of *A. thaliana* by measuring macroscopic disease symptoms, size differences, and molecular markers of disease. For the field collected plants in this study, it was unclear how the colonization levels we observed related to microbial colonization observed in the laboratory. We therefore wanted to make use of information from laboratory experiments to interpret the field data. Having obtained information about the progression of infections with *P. syringae* that is either recognized by the host immune system (incompatible) or unrecognized (compatible), we were in a position to interpret the loads we had observed in the field. More specifically, we expected that this information would provide a baseline for distinguishing between compatible and incompatible interactions in the field.

In the laboratory, *P. syringae* abundance increased over the first three days after infection, with mean infection levels of compatible infections on day 2 exceeding those of incompatible infections on any day. We thus considered the mean load achieved by DC3000:EV on day 2 to exceed the “resistance threshold” of the infection. In the field, we found that 6% of plants contained a Pseudomonadaceae load that exceeded the maximum infection level in incompatible infections. The second most common bacterial family in Germany comprises the Sphingomonadaceae, but there is no evidence in the literature that they can be pathogens, and their load did not exceed the resistance threshold in any of the plants analyzed (Figure 4).

**Figure 4:**
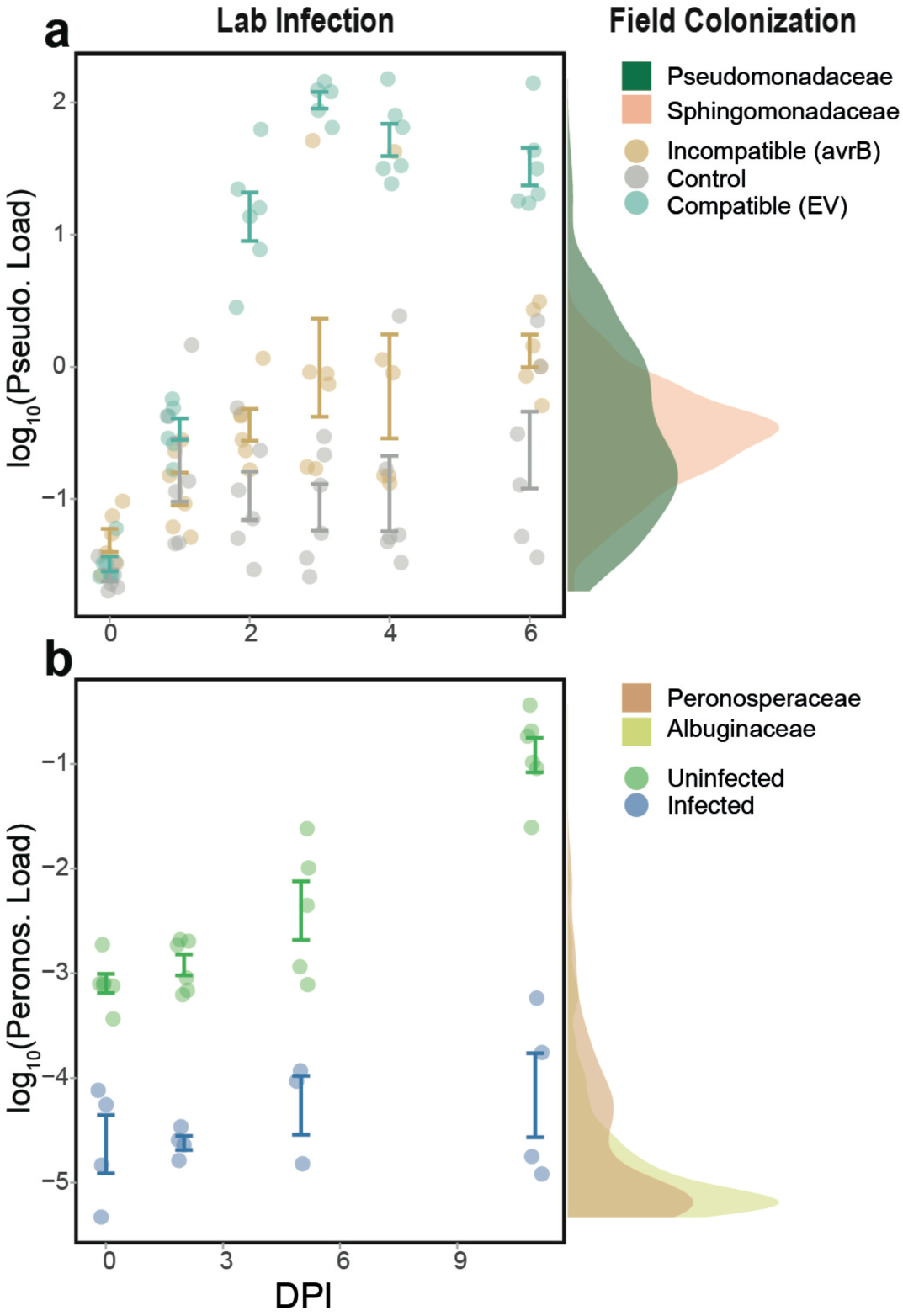
Relating metagenomic load in the lab to that in the field. The ratio of microbial genome coverage to *A. thaliana* genome coverage was compared between the controlled infections and the field data. (a) Comparison of the data from Pst DC3000 infections in the laboratory and the loads observed for two of the most abundant bacterial families—Pseudomonadaceae and Sphingomonadaceae. The left panel shows the load in the laboratory (same data as in figure 3 but with more observations), and the right shows a density plot of the load assessed in the field populations. The y-axis is shared between left and right panels. (b) Comparison of the data from *HpA* infections in the laboratory and the loads observed for two of the most abundant Oomycete families– Peronosporaceae and Albuginaceae.

As illustrated in Figures 3 and 4 (for Pseudomonadaceae), in the bacterial microbiome from the field, only Pseudomonadaceae and Burkholderiaceae–a family containing pathogenic members such as the genus *Ralstonia* [43]–colonized plants to a level that surpassed the mean colonization level of the incompatible *P. syringae* interaction (DC3000:avrB) in the lab. Similarly, only Peronosporaceae and Albuginaceae achieved a load that exceeded the day 2 resistance threshold for oomycetes, when hyphae and spores were still rarely observed in samples. The remaining taxa in the microbiota of wild plants were not found above levels of laboratory plants infected with microbes that did not induce an immune response. From these results alone it is not yet clear whether the plants recognize the abundance of low-abundance microbes and suppress growth, or suppress growth through cross-resistance.

Note that while several of the same bacterial and oomycete families were found to contribute to microbial load across populations, it is likely that in different regions and plants, the actual species or strains were likely not the same, since strains of species, as well as their gene content and pathogenic capacity, are known to differ within and between populations [44].

## Discussion

Hundreds of microbial families colonize *A. thaliana* leaves, and the colonizing microbiota differs between plant genotypes [45, 46] and regions [5, 46]. In this study we found that very few of these microbial families–not even 1 in a 100–ever proliferate to large population sizes i*n planta*, with fewer than 10% of plants being extensively colonized to levels equivalent to late-stage microbial infections in the laboratory.

While composition and total load differed across regions, the identity of microbial families that successfully proliferated *in planta* was largely the same across *A. thaliana* populations. These exceptional families–Pseudomonadaceae, Peronosporaceae, and Albuginaceae–contained known *A. thaliana* pathogens that proliferated in both regions to loads higher than the average achieved in incompatible infections in the laboratory. This extensive proliferation occurred only in a minority of plants, and we hypothesize that these plants were undergoing pathogenic infection. Importantly, we find that high load of a taxon was associated with two leaf microbiome summary statistics: differences in composition (Figure 2) and reductions in alpha diversity (Figure 3). The relationship between the presence of a single prolific taxon and microbial diversity is so clear in our dataset that it should be possible to take low microbial diversity in the phyllosphere as an initial classifier for latent disease both in wild and cultivated plants. Follow-up work could examine the host for signs of compromised health and test the disease capacity of the identified microbe across a range of environments.

There are several explanations for the limited loads achieved by “background taxa” in our dataset. One possibility is that the background taxa are controlled by the plant immune system and it is only when a microbe successfully evades [47, 48] or overcomes [49] the immune system that it can proliferate beyond the level of incompatible infection resistance. Whether these microbes cannot proliferate successfully *in planta* under any conditions, or are instead controlled by the plant immune system, remains an open question. In one example, Kniskern and colleagues [50] demonstrated that immune-deficient *A. thaliana* mutants planted in the field exhibit increased microbial loads, indicating host control of the larger microbial community. We now have the tools to probe this question with greater specificity to determine on a strain-specific level which microbes are controlled by the host immune system.

There is a growing body of evidence that commensal microbes interact with the plant immune system. For example, specific strains of *Sphingomonas*, an abundant, putatively commensal genus in the phyllosphere, stimulate plant immune responses [1], and one can envision that their proliferation is kept in-check by elements of the plant immune system that also control *bona fide* pathogens [51, 52]. In addition, many microbes can probably not easily proliferate in the nutrient-limited conditions of the phyllosphere. While this possibility has not been explicitly tested, in the root system it is well established through stable isotope profiling that only a fraction of the microbiome measurably proliferates at any point in time [53]. Lastly, there is a possibility that unknown biases in our extraction and sequencing methods could potentially skew the relative abundance of microbiota. While this is true of every extraction and sequencing method to date, metagenomic results are concordant with those of the 16S rDNA data [16], and the prominence of Pseudomonadaceae, Peronosporaceae, Sphingomonadaceae, and Albuginaceae in wild phyllospheres has been observed repeatedly across experiments and methodologies [2, 3, 45, 54].

Our findings of low microbial load and a negative relationship between load and alpha diversity point to a paradigm in the leaf apoplast that differs from that in the human gut and perhaps that in the root. While a healthy human gut is colonized by large numbers of microbes interacting symbiotically with the host, and lower bacterial levels can even be associated with non-microbial disease states [7], our results suggest that a healthy leaf apoplast likely does not house high numbers of microbes.

Unlike in the human gastrointestinal tract or in the plant root system, there are only a few examples of *bona fide* commensals inside the plant leaf. The plant apoplast is a nutrient-limited intercellular space [55] used by plants for respiration. Within the leaves, few microbes serve useful functions in these basic cellular processes [56], and instead most are known to interfere with them [57]. The few cases in the leaf apoplast in which microbes have been found to be beneficial are largely due to their ability to protect the plant against pathogens [1] (but see the work by Mayak and colleagues [58], who describe what may be an endophytic association). It seems plausible that it is only when a pathogenic microbe successfully evades the host immune responses that it can colonize to high titer in the phyllosphere.

Knowledge of microbial load may be particularly informative when one wants to identify microbes with an effect on plant fitness. There is an increasing appreciation of the role of microbe-derived small molecules in interactions with the plant [59]. Many of these small molecules are regulated by quorum sensing mechanisms [60, 61], which signal a certain population size of the focal microbe. Indeed, achieving appreciable metabolite levels to influence host function may require high microbial load. Furthermore, because few microbes in the plant apoplast benefit the host, a reasonable hypothesis going forward is that a high microbial load is a signature of pathogenic (or pre-pathogenic) infection. We conclude that the methods we have developed and implemented can be used not only to monitor plants for pathogenic infection, but also to discover novel disease-causing microbes.

## Methods

### Field sample collections

Metagenomic samples from Germany were previously described and published [2]. Briefly, these samples were collected from four populations in Southwestern Germany over two seasons. Metagenomic samples from Sweden were collected from Southern Sweden in March 2017 and from Northern Sweden in April 2017. The locations and dates of collection for all samples are included in Table S1. Whole rosettes were collected from the field with tweezers and scissors sterilized between plant samples, washed in sterile water, then flash frozen on dry ice. The samples were then stored at -80°C until time of DNA extraction.

### qPCR quantification

To assess bacterial abundance in infections, we amplified a fragment of bacterial 16S rDNA using a forward primer that begins priming at position 799 of the 16S rDNA locus [62], and a reverse primer that begins priming at position 902 [63]. (799F-AACMGGATTAGATACCCKG and 902R-GTCAATTCMTTTGAGTTTYARYC). 902R was designed specifically for the exclusion of chloroplast (and cyanobacterial) amplification [63]. Assessment of mitochondrial and chloroplast amplification was made by in silico PCR (with blast) against NC_037304.1(mitochondrion) and NC_000932.1 (chloroplast). To assess plant abundance in the samples, we amplified a segment of the *A. thaliana* single-copy gene *GIGANTEA* (F-ACATGCTTTGATACAGCGGTGA and R-TGGATTCATTTCAGTCCTTGAGG). qPCR was done on a BioRad CFX384 Real-time System and analyzed with the CFX Manager Software. The following conditions were used for amplification of

16S rDNA:

1. 95°C for 3 minutes
2. 95°C 15 seconds
3. 53°C 30 seconds
4. 72°C 30 seconds
5. Return to (2) 44 times
6. Melting curve

*GIGANTEA* (*GI*) :

(1) 95°C for 3 minutes
(2) 95°C 10 seconds
(3) 55°C 30 seconds
(5) Return to (2) 45 times
(6) Melting curve

The standard curve for 16S rDNA amplification was established with serial dilutions of a pure extraction of Pst DC3000. Serial dilutions of Col-0 DNA were used to establish a standard curve for *GI* amplification.

### Metagenomic library analysis

Total DNA was extracted from rosettes that had been flash-frozen in liquid N_2_ by grinding plant tissue in micro-tubes filled with garnet rocks. Further metagenomic DNA purification and library preparation protocol was as published [2]. Library molecules were size selected on a Blue Pippin instrument for the size range 350-750 base pairs (Sage Science, Beverly, MA, USA). Multiplexed libraries were sequenced with 2×150 base pairs paired-end reads on an HiSeq3000 instrument (Illumina).

A significant challenge in the analysis of plant metagenomic sequences is the proper masking of the host DNA. In order to mask host derived sequences, reads were mapped against the *A. thaliana* TAIR10 Col-0 reference genome [64] with *bwa mem* [34] using standard parameters. We have shown that metagenomic load can be reliably assessed with as few as 30,000 reads [16]. After removing samples that had fewer than 30,000 reads, all read pairs flagged as not mapping to *A. thaliana* were extracted with *samtools* [65] as the putatively “metagenomic” fraction. Average coverage of the *A. thaliana* genome was assessed by the *samtools* ‘depth’ command, taking the average over the five nuclear chromosomes (excluding mitochondria and chloroplast DNA). The remaining unmapped reads were then classified using the metagenomic classification tool *centrifuge* [35] (standard parameters). Taxonomic assignment at the family level was considered for downstream analyses.

The output read table from centrifuge was normalized using custom scripts (https://tkarasov.github.io/controlled_metagenomics) to assess for each sample the average coverage per each microbial family/the *A. thaliana* genome. Average genome size of a microbial family was estimated using information downloaded from https://www.ncbi.nlm.nih.gov/genome/browse/#!/overview/ on 9/21/2018 and an NCBI taxonomy generated by the ‘tools/taxdmp2tree’ script in the Ultimate edition of MEGAN [66] in September 2018. For families for which genome size data was not available, a default size of 3.87 Mb (the average bacterial genome size) was assigned for bacteria [67], 8.97 Mb for fungi [68], and 37 Mb for oomycetes [69]. Note that the assigned values for fungi and oomycete are the lower bounds for genome sizes for these taxa, hence this assignment is likely to inflate the estimate of abundance of genomes with less complete assemblies (i.e., typically less well-studied organisms). If a genome assembly was smaller than 1 Mb, we deemed the assembly to be unreliable and assigned the same missing value for size. Less than 3% of families detected in *centrifuge* assignment had no genome size estimate. Note that all subsequent comparisons are based off of the ratio of microbial/plant abundance.

To compare composition and abundance between regions we acknowledged differences in sequencing depth between samples by transforming the genome coverage data to variance-stabilized estimates with the R package DESeq2 [37, 70].

Because standard compositional comparisons such as those in 16S rDNA amplicon sequencing are constrained by their compositional nature, we compare ratios (with plant coverage as the denominator), which avoids bias [18] that can arise in compositional comparisons, including erroneous conclusions regarding correlations between taxa[71]. All scripts for metagenome mapping and figures can be found at: https://tkarasov.github.io/controlled_metagenomics. Diversity analyses were performed in R (version 3.5.3) using both custom scripts and the *vegan* package [72]. Because the sampling was unbalanced (many more samples from specific populations in Germany than in Sweden), we repeatedly subsampled (n = 100 bootstraps) eight individuals per population for Principal Coordinate Analysis and multivariate regression. The values for variance explained are the means over these 100 bootstraps. Metagenomic reads from Swedish populations can be accessed in European Nucleotide Archive via PRJEB34580. Metagenomic reads from German populations can be accessed at the European Nucleotide Archive via Primary Accession PRJEB24450.

### *Pseudomonas syringae* infections

Seeds of the *A. thaliana* genotype Col-0 were frozen overnight at -80°C and bleach sterilized. The sterilized seeds were then planted on soil and grown with 8-hours of light per day at 23°C with a high humidity dome. At 42 days of age, the plants were infected with DC3000:EV, DC3000:avrB or control (10 mM MgSO_4_) as described in the paragraph below. Plants were labeled and randomized across three flats.

*P. syringae* strain Pst DC3000 was used for all infections. The strain used for infections was transformed with either the plasmid pMH223 encoding kanamycin (KM) resistance only or pMH221:AvrB. DC3000 was first made electrocompetent via sucrose washes [73] and transformed with the constructs described. Successful transformants were selected on plates containing 50 µg/mL kanamycin. The night prior to infection, a colony was inoculated into 1-5 mL of Luria broth (LB) containing 10 µg/mL KM, and the culture was grown overnight at 28°C. In the morning, the sample was diluted 1:10 in 5 mL of fresh LB and grown for 2 to 4 hours. The resulting culture was spun down at 3500 x g, and resuspended in 10mM MgSO_4_ and diluted to an OD_600_ of 0.0002. Four leaves were syringe-inoculated per plant. Six replicates per treatment per day were taken at five timepoints: the first was taken immediately upon infection, the next after one day, then two days, three days, four days and a final time-point six days post infection. For collection of infected or uninfected material, sterile forceps were used to remove four infected leaves per plant which were then flash-frozen immediately in liquid nitrogen and stored at -80°C. Leaves used for CFU counting were surface-sterilized in 70% EtOH for five seconds. Two hole punches per two sterilized leaves were taken using gelatin capsules (Kapselwelt product number 1002).

### *HpA* infections

Seeds of the *A. thaliana* genotype H2081 belonging to the HPG1 haplogroup [74] were frozen overnight at -80°C and bleach sterilized. This genotype was chosen for its susceptibility to the focal *HpA* genotype. The sterilized seeds were then planted on soil and grown with 8-hours of light per day at 23°C with a high humidity dome. After 6 weeks plants were moved to a Percival growth chamber with ten-hours of light per day and a cooler temperature of 15°C. At 42 days of age, the plants were infected with *HpA* genotype 140HML004, a genotype we knew to be able to successfully colonize *A. thalian*a H2081. For infection, sporangiospores of 140HML004 propagated in Ws-0 *eds1 (eds1-1)* mutants [75] were collected and diluted in double distilled H_2_O to a concentration of 54,000 spores/mL. Three 5 µL drops of this spore solution were inoculated on either side of the mid-vein of five leaves per plant. This totaled 60 µl of spore solution per leaf, or an estimated 3,240 spores per leaf. Control plants were inoculated with the same procedure but with ddH_2_O instead of spore solution. Plants were labeled and randomized by day within a flat of *HpA* treatment or control treatment. For collection of infected or uninfected material, sterile forceps were used to remove four infected leaves per plant which were then flash-frozen immediately in liquid nitrogen and stored at -80°C.

Progression of *HpA* infections was monitored using Trypan blue staining [13, 29]. Briefly, Trypan blue is known to stain the external structures of *HpA* hyphae [13]. Leaves were removed from the rosette, heated for one hour at 70°C in Trypan blue stain solution (10 ml ddH_2_0, 10 ml phenol, 10 ml lactic acid, 10 ml glycerol, and 20 mg Trypan blue and water mixed in 1:2 ratio with 95% ethanol). Decolorization of stained leaves was achieved via soaking in aqueous chloral hydrate (2.5 g of chloral hydrate per 1ml ddH_2_0) until leaves were largely translucent (from two days to two weeks). Leaves were then mounted in 60% glycerol for observation on a Zeiss AxioImager Z1.

### Comparison of bacterial and plant growth via luminescence

Plant genotypes Eyach 15-2 and Col-0 were used. Seeds were stratified for twelve days at 4°C and then were grown in long-day (16 h) at 23°C. After 5 days, seedlings were transferred to 24-well plates with ½ strength MS medium (Duchefa M0255.0050) and 1% agar, one seedling per well. Six days after, i.e. when plants were 11-days old, they were infected with single bacterial strains.

*Pseudomonas* strains were transformed to express the *lux* operon via electroporation. pUC18-mini-Tn7T-Gm-lux was a gift from Herbert Schweizer (Addgene plasmid # 64963). *Pseudomonas* strains were grown overnight at 28°C in Luria-Bertani (LB) medium with 100 ng/mL of nitrofurantoin, diluted the following morning 1:10 in 5 mL selective medium and grown for 3 additional hours. Bacteria were then pelleted at 3,500 g and brought to an OD_600_ of 0.01 in 10 mM MgSO_4_. 100 µL of this bacterial suspension were used to drip-inoculate plants, distributing the volume over the whole rosette. Plants were mock infected with 10 mM MgSO_4_ as control. Plates were returned to the growth chamber, and three days after infection whole rosettes were cut for luminescence quantification.

For luminescence quantification whole rosettes were transferred to 96 deep-well plates (2.2 mL, Axygen), containing two 5 ± 0.03 mm glass beads (Roth) and 400 µL of 10 mM MgSO_4_, and ground for one minute at 20 m/s in a TissueLyser II (QIAGEN). Then, 10 mM MgSO_4_ was added to a final volume of 1 mL, and 200 µL were transferred to a 96-well Lumitrac white plate. Luminescence was measured in a multiplate reader (TECAN Infinite F200) with 2,000 ms of integration time. Each well was measured three times, and the mean was calculated for further analysis. The signal of MgSO_4_ blanks was subtracted from the samples’ signal before analysis.

Data analysis was conducted in R 3.5.1. Luminescence signal was log-transformed, and the NA generated due to negative luminescence values (obtained after subtraction of blanks) were replaced by 0, to retain the meaning of absence of luminescence.

### Image analysis

For plant growth quantification, plates were photographed before infection and seven days post infection, with a tripod-mounted Canon PowerShot G12 digital camera. Individual plants were extracted from whole-plate images. The number of green pixels was determined for each plant and used as a proxy for plant fresh mass. The segmentation of the plant from background was performed by applying thresholds in Lab color space, followed by a series of morphological operations to remove noise and non-plant objects. Finally, a GrabCut-based postprocessing was applied and csv files with plant IDs and green pixel counts were created. The workflow was implemented in Python 3.6 and bash using OpenCV 3.1.0 and scikit-image 0.13.0 for image processing operations.

### Cryogenic electron microscopy

*A. thaliana* plants Col-0 were syringe infected with Pst DC3000 (OD_600_=0.0002) as described above after 4 weeks of growing in soil. The leaf material was collected three days after infection and immediately fixed with 2.5% glutaraldehyde in PBS-buffer. After 1-hour incubation at room temperature (approximately 25°C), the samples were transferred to 4°C for three days. Subsequently, four washes in PBS buffer were carried out over two days. Immediately prior to visualization, samples were carefully dried with paper tissues and compressed into a metal holder. Freezing was performed inside a cryo loading chamber filled with liquid N_2_. Frozen leaves were shattered with a metal tool tapped onto the edges. Sample transfers were conducted with the Vacuum cryo transfer system VCT100.

All samples were imaged on a Zeiss LEO Crossbeam 1540 Scanning Electron Microscope with a VCT100 cryo load lock system. The samples were sublimated inside the microscope by increasing the temperature from -140°C to 90°C. Afterwards the samples were sputter coated with platinum in a cryo sputter coater Baltec SCD500. The image was recorded with an Everhart Thornley Detector at 114 X magnification and 3 kV beam voltage at -140°C temperature.

## Acknowledgements

We thank members of the Swedish collection team for participating in the collecting of the *A. thaliana* samples in Sweden. We thank Michael Werner, Hernán Burbano, Rebecca Satterwhite and Andrew Gloss for comments on the manuscript.

## Funding

Funding was provided by HFSP Long-Term Fellowships (TLK, DSL), an EMBO Long-Term Fellowship (TLK), NIH R01 GM 083068 (JB), ERC Advanced Grant IMMUNEMESIS (340602), the DFG through SPP Priority Program DECRyPT, the Max Planck Society (DW).

## Author Contributions

TLK and DW devised the study. TLK, MN, SK, BS and ADJ performed the experiments, TLK and MN analyzed the data. IB contributed scripts for image analysis. ES and GS advised on *HpA* infections. JR and DSL advised on metagenomic analysis. JB provided samples from Sweden. TLK and DW wrote the manuscript with help from all authors.

## Competing Interests

The authors declare no competing interests.

---

**Figure S1:**
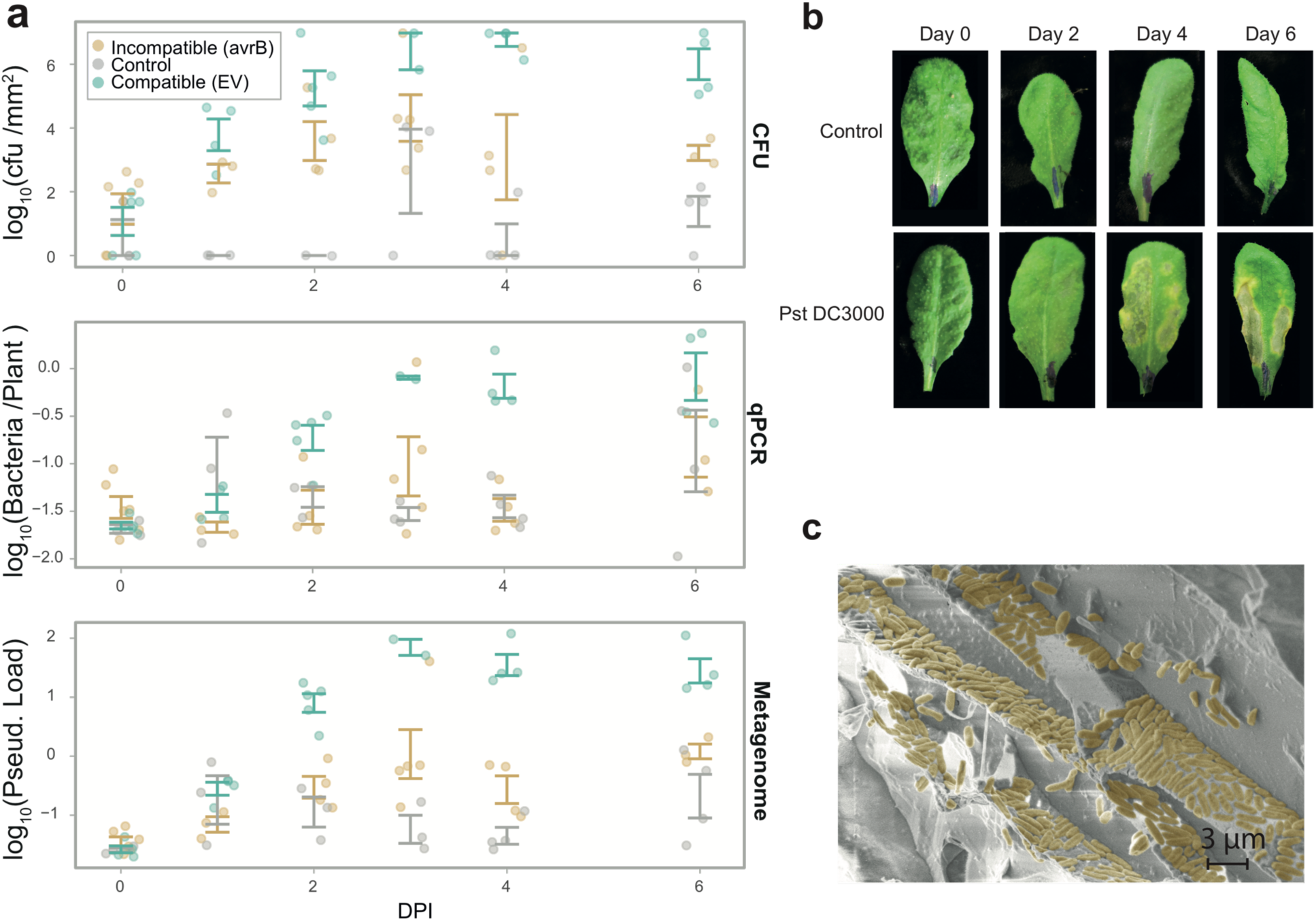
Comparison of different metrics of *P. syringae* growth *in planta*. (a) Time-series of *P. syringae* growth (measured in colony forming units [cfu] per mm2), metagenomic load of Pseudomonadaceae, and qPCR measurement of bacterial 16S rDNA abundance. Data is presented as mean values +/- standard errors. (b) Macroscopic image of infection with Pst DC3000 in a time-series. (c) Scanning electron microscopic image of Pst DC3000 infection in Col-0 at three days post infection. Related to Figure 1.

**Figure S2:**
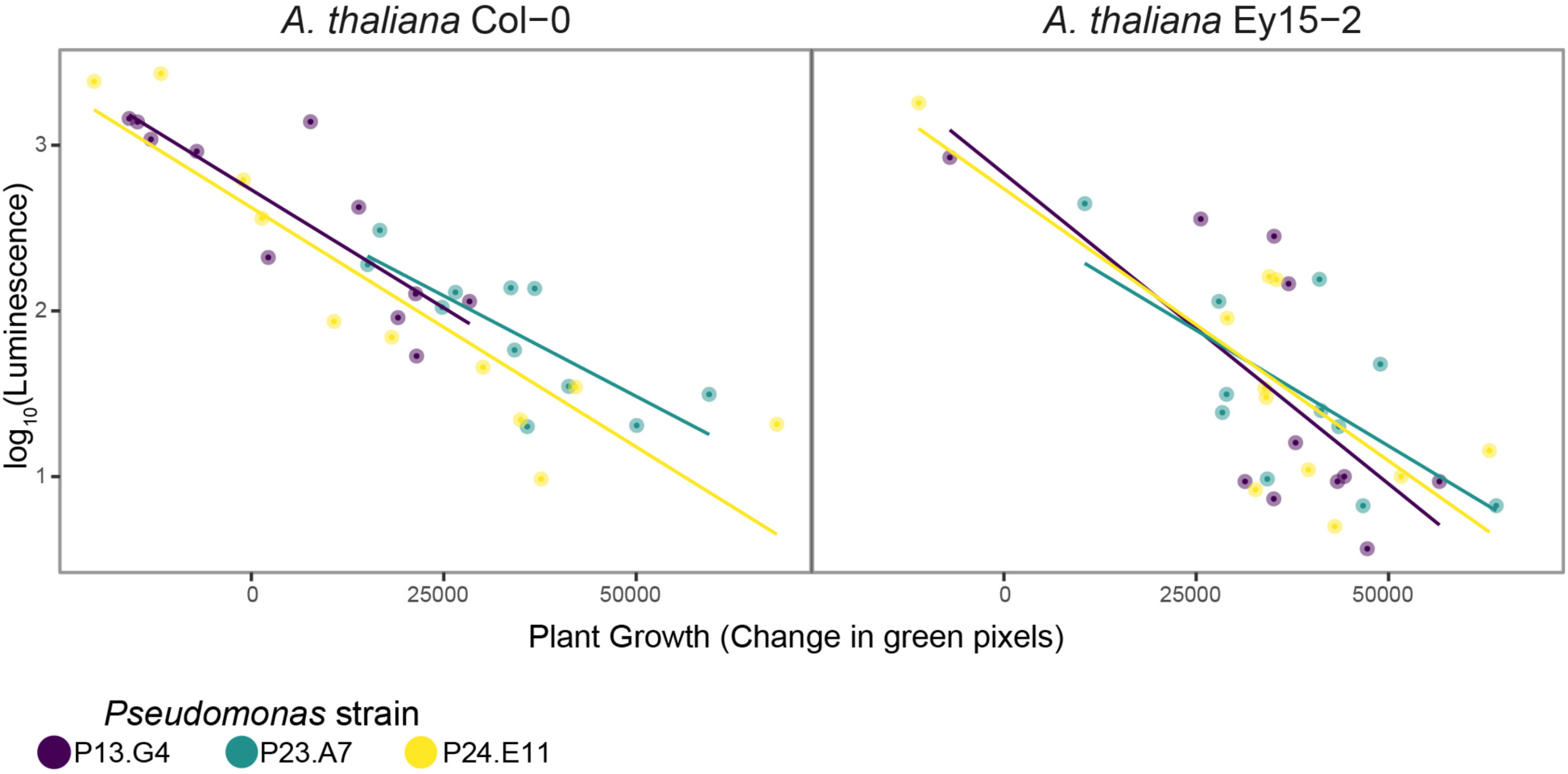
*Pseudomonas* growth is associated with reduced *A. thaliana* growth. The growth of three *Pseudomonas* isolates (Karasov et al. 2018) and two *A. thaliana* genotypes were measured simultaneously in laboratory infections. Increased *Pseudomonas* growth is significantly associated with reduced *A. thaliana* growth across all *Pseudomonas* and *A. thaliana* genotypes. The graph shows the relationship between bacterial growth (measured in luminescence) and plant growth (measured in green pixels from image analysis). The lines represents the line of best fit for each individual strain from linear regression. Related to Figure 2.

**Figure S3:**
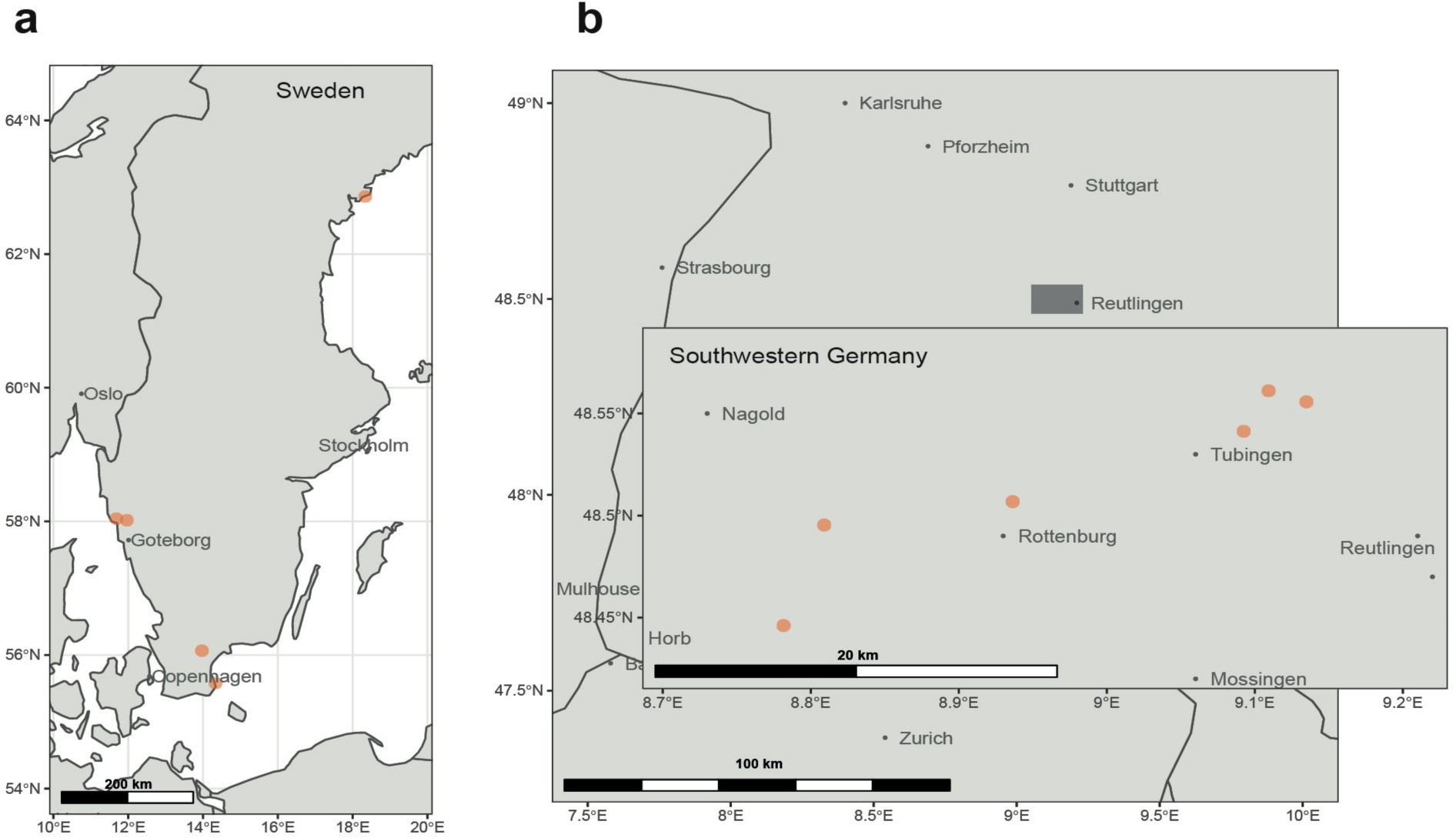
Maps of collections locations in Germany and Sweden. Plants were collected in Germany and sequenced as described (Karasov et al. 2018). Plants were collected in Sweden in this study and the same procedure was followed as for the German plants for metagenomic extraction and sequencing. Related to Figure 2.

**Figure S4:**
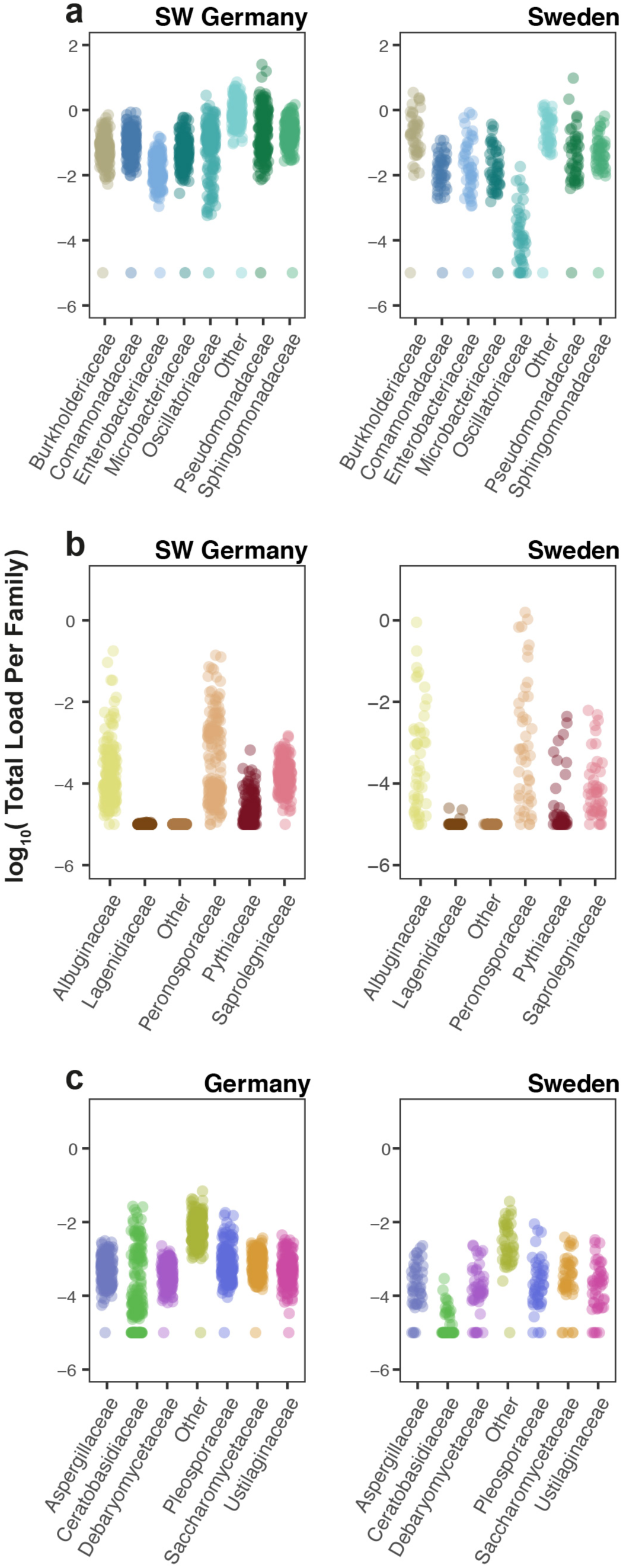
The distribution of microbial load per family. Categorical plots showing the distribution of loads per family across plants in Germany and Sweden. All panels share the same y-axis label. (a) Bacterial families (b) Oomycete families and (c) Fungal families. The category “Other” represents the sum of all remaining taxa that are not shown.

**Figure S5:**
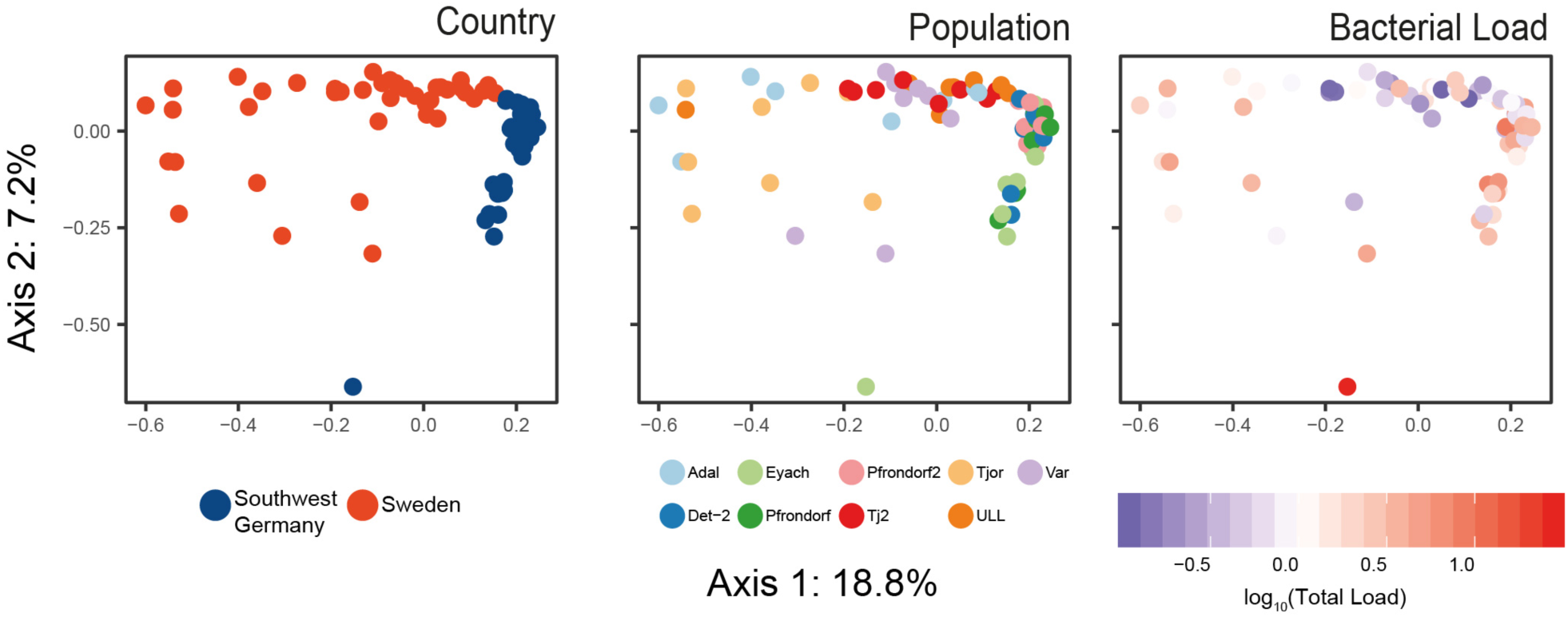
The relationship between location, load and microbiome composition. PCoA on Bray-Curtis dissimilarity matrices calculated from a balanced sampling from populations of the relative abundance of the bacteria included in Figure 1a. Panels are colored according to region, according to population (nested in region), and according to load.

**Figure S6:**
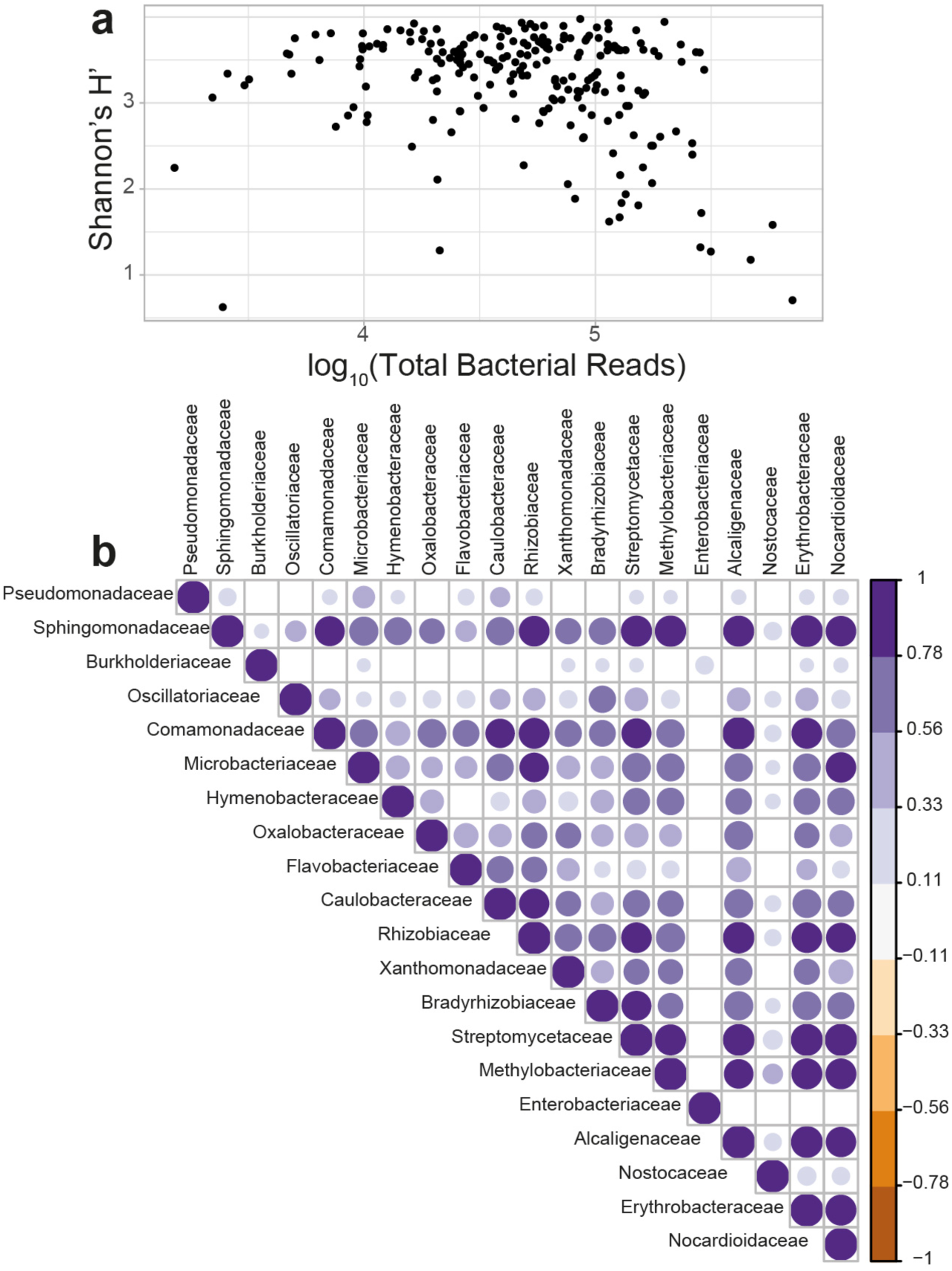
High load is associated with proliferation of single taxa rather than suppression of others. (a) Scatterplot showing the relationship between the number of reads assigned as bacterial in centrifuge (Kim et al. 2016) and the estimated Shannon Diversity Index (H’). (b) Correlogram showing significant correlations (p < 0.05, after correction for multiple testing) in the abundance of the top 20 most abundant bacterial families. Related to Figure 3.

## Notes

#### Summary of Updates

Reordered presentation

## References

1. Innerebner G, Knief C, Vorholt JA. Protection of Arabidopsis thaliana against leaf-pathogenic Pseudomonas syringae by Sphingomonas strains in a controlled model system. Appl Environ Microbiol 2011; 77: 3202–3210.

2. Karasov TL, Almario J, Friedemann C, Ding W, Giolai M, Heavens D, et al. Arabidopsis thaliana and Pseudomonas Pathogens Exhibit Stable Associations over Evolutionary Timescales. Cell Host Microbe 2018; 24: 168–179.

3. Lundberg DS, Lebeis SL, Paredes SH, Yourstone S, Gehring J, Malfatti S, et al. Defining the core Arabidopsis thaliana root microbiome. Nature 2012; 488: 86–90.

4. Kembel SW, O’Connor TK, Arnold HK, Hubbell SP, Wright SJ, Green JL. Relationships between phyllosphere bacterial communities and plant functional traits in a neotropical forest. Proc Natl Acad Sci U S A 2014; 111: 13715–13720.

5. Wagner MR, Lundberg DS, Del Rio TG, Tringe SG, Dangl JL, Mitchell-Olds T. Host genotype and age shape the leaf and root microbiomes of a wild perennial plant. Nat Commun 2016; 7: 12151.

6. Vaughn DW, Green S, Kalayanarooj S, Innis BL, Nimmannitya S, Suntayakorn S, et al. Dengue viremia titer, antibody response pattern, and virus serotype correlate with disease severity. J Infect Dis 2000; 181: 2–9.

7. Vandeputte D, Kathagen G, D’hoe K, Vieira-Silva S, Valles-Colomer M, Sabino J, et al. Quantitative microbiome profiling links gut community variation to microbial load. Nature 2017; 551: 507–511.

8. Goodrich JK, Waters JL, Poole AC, Sutter JL, Koren O, Blekhman R, et al. Human genetics shape the gut microbiome. Cell 2014; 159: 789–799.

9. Maignien L, DeForce EA, Chafee ME, Eren AM, Simmons SL. Ecological succession and stochastic variation in the assembly of Arabidopsis thaliana phyllosphere communities. mBio 2014; 5: e00682–13.

10. Jakob K, Goss EM, Araki H, Van T, Kreitman M, Bergelson J. Pseudomonas viridiflava and P. syringae—natural pathogens of Arabidopsis thaliana. Mol Plant Microbe Interact 2002; 15: 1195–1203.

11. Coates ME, Beynon JL. Hyaloperonospora arabidopsidis as a pathogen model. Annu Rev Phytopathol 2010; 48: 329–345.

12. Greenberg JT, Guo A, Klessig DF, Ausubel FM. Programmed cell death in plants: a pathogen-triggered response activated coordinately with multiple defense functions. Cell 1994; 77: 551–563.

13. Van Damme M, Andel A, Huibers RP, Panstruga R, Weisbeek PJ, Van den Ackerveken G. Identification of Arabidopsis loci required for susceptibility to the downy mildew pathogen Hyaloperonospora parasitica. Mol Plant Microbe Interact 2005; 18: 583–592.

14. Stämmler F, Gläsner J, Hiergeist A, Holler E, Weber D, Oefner PJ, et al. Adjusting microbiome profiles for differences in microbial load by spike-in bacteria. Microbiome 2016; 4: 28.

15. Kleiner M, Thorson E, Sharp CE, Dong X, Liu D, Li C, et al. Assessing species biomass contributions in microbial communities via metaproteomics. Nat Commun 2017; 8: 1558.

16. Regalado J, Lundberg DS, Deusch O, Kersten S, Karasov T, Poersch K, et al. Combining whole genome shotgun sequencing and rDNA amplicon analyses to improve detection of microbe-microbe interaction networks in plant leaves. bioRxiv. 2019., 823492

17. Wu GD, Lewis JD, Hoffmann C, Chen Y-Y, Knight R, Bittinger K, et al. Sampling and pyrosequencing methods for characterizing bacterial communities in the human gut using 16S sequence tags. BMC Microbiol 2010; 10: 206.

18. McLaren MR, Willis AD, Callahan BJ. Consistent and correctable bias in metagenomic sequencing measurements. bioRxiv. 2019., 559831

19. Suzuki MT, Giovannoni SJ. Bias caused by template annealing in the amplification of mixtures of 16S rRNA genes by PCR. Appl Environ Microbiol 1996; 62: 625–630.

20. Ziesemer KA, Mann AE, Sankaranarayanan K, Schroeder H, Ozga AT, Brandt BW, et al. Intrinsic challenges in ancient microbiome reconstruction using 16S rRNA gene amplification. Sci Rep 2015; 5: 16498.

21. Wanner LA, Mittal S, Davis KR. Recognition of the avirulence gene avrB from Pseudomonas syringae pv. glycinea by Arabidopsis thaliana. Mol Plant Microbe Interact 1993; 6: 582–591.

22. Velásquez AC, Oney M, Huot B, Xu S, He SY. Diverse mechanisms of resistance to Pseudomonas syringae in a thousand natural accessions of Arabidopsis thaliana. New Phytol 2017; 214: 1673–1687.

23. Innes RW, Bisgrove SR, Smith NM, Bent AF, Staskawicz BJ, Liu YC. Identification of a disease resistance locus in Arabidopsis that is functionally homologous to the RPG1 locus of soybean. Plant J 1993; 4: 813–820.

24. Pel MJC, van Dijken AJH, Bardoel BW, Seidl MF, van der Ent S, van Strijp JAG, et al. Pseudomonas syringae evades host immunity by degrading flagellin monomers with alkaline protease AprA. Mol Plant Microbe Interact 2014; 27: 603–610.

25. Cuppels DA. Generation and Characterization of Tn5 Insertion Mutations in Pseudomonas syringae pv. tomato. Appl Environ Microbiol 1986; 51: 323–327.

26. Kniskern JM, Barrett LG, Bergelson J. Maladaptation in wild populations of the generalist plant pathogen Pseudomonas syringae. Evolution 2011; 65: 818–830.

27. Koch E, Slusarenko A. Arabidopsis is susceptible to infection by a downy mildew fungus. Plant Cell 1990; 2: 437–445.

28. Hardham AR. Cell biology of plant-oomycete interactions. Cell Microbiol 2007; 9: 31–39.

29. Strober W. Trypan blue exclusion test of cell viability. Curr Protoc Immunol 2001; Appendix 3: Appendix 3B.

30. Gao L, Roux F, Bergelson J. Quantitative fitness effects of infection in a gene-for-gene system. New Phytol 2009; 184: 485–494.

31. Roux F, Gao L, Bergelson J. Impact of initial pathogen density on resistance and tolerance in a polymorphic disease resistance gene system in Arabidopsis thaliana. Genetics 2010; 185: 283–291.

32. Dunn OJ. Multiple Comparisons among Means. J Am Stat Assoc 1961; 56: 52–64.

33. Thiergart T, Duran P, Ellis T, Garrido-Oter R, Kemen E. Root microbiota assembly and adaptive differentiation among European Arabidopsis populations. bioRxiv 2019.

34. Li H. Aligning sequence reads, clone sequences and assembly contigs with BWA-MEM. arXiv [q-bioGN]. 2013.

35. Kim D, Song L, Breitwieser FP, Salzberg SL. Centrifuge: rapid and sensitive classification of metagenomic sequences. Genome Res 2016; 26: 1721–1729.

36. McMurdie PJ, Holmes S. phyloseq: an R package for reproducible interactive analysis and graphics of microbiome census data. PLoS One 2013; 8: e61217.

37. McMurdie PJ, Holmes S. Waste not, want not: why rarefying microbiome data is inadmissible. PLoS Comput Biol 2014; 10: e1003531.

38. McArdle BH, Anderson MJ. Fitting multivariate models to community data: a comment on distance-based redundancy analysis. Ecology 2001; 82: 290–297.

39. Benjamini Y, Hochberg Y. Controlling the False Discovery Rate: A Practical and Powerful Approach to Multiple Testing. J R Stat Soc Series B Stat Methodol 1995; 57: 289–300.

40. Hatfield JL, Prueger JH. Temperature extremes: Effect on plant growth and development. Weather and Climate Extremes 2015; 10: 4–10.

41. Woo NS, Badger MR, Pogson BJ. A rapid, non-invasive procedure for quantitative assessment of drought survival using chlorophyll fluorescence. Plant Methods 2008; 4: 27.

42. Lipiec J, Doussan C, Nosalewicz A, Kondracka K. Effect of drought and heat stresses on plant growth and yield: a review. International Agrophysics 2013; 27: 463–477.

43. Denny T. Plant pathogenic Ralstonia species. In: Gnanamanickam SS (ed). Plant-Associated Bacteria. 2006. Springer Netherlands, Dordrecht, pp 573–644.

44. Karasov TL, Barrett L, Hershberg R, Bergelson J. Similar levels of gene content variation observed for Pseudomonas syringae populations extracted from single and multiple host species. PLoS One 2017; 12: e0184195.

45. Horton MW, Bodenhausen N, Beilsmith K, Meng D, Muegge BD, Subramanian S, et al. Genome-wide association study of Arabidopsis thaliana leaf microbial community. Nat Commun 2014; 5: 5320.

46. Brachi B, Filiault D, Darme P, Le Mentec M, Kerdaffrec E, Rabanal F, et al. Plant genes influence microbial hubs that shape beneficial leaf communities. bioRxiv. 2017., 181198

47. Glazebrook J. Contrasting mechanisms of defense against biotrophic and necrotrophic pathogens. Annu Rev Phytopathol 2005; 43: 205–227.

48. Okmen B, Doehlemann G. Inside plant: biotrophic strategies to modulate host immunity and metabolism. Curr Opin Plant Biol 2014; 20: 19–25.

49. van Kan JAL. Licensed to kill: the lifestyle of a necrotrophic plant pathogen. Trends Plant Sci 2006; 11: 247–253.

50. Kniskern JM, Traw MB, Bergelson J. Salicylic acid and jasmonic acid signaling defense pathways reduce natural bacterial diversity on Arabidopsis thaliana. Mol Plant Microbe Interact 2007; 20: 1512–1522.

51. Hacquard S, Spaepen S, Garrido-Oter R, Schulze-Lefert P. Interplay Between Innate Immunity and the Plant Microbiota. Annu Rev Phytopathol 2017; 55: 565–589.

52. Vannier N, Agler M, Hacquard S. Microbiota-mediated disease resistance in plants. PLoS Pathog 2019; 15: e1007740.

53. Gkarmiri K, Mahmood S, Ekblad A, Alström S, Högberg N, Finlay R. Identifying the Active Microbiome Associated with Roots and Rhizosphere Soil of Oilseed Rape. Appl Environ Microbiol 2017; 83: 01938–01917.

54. Agler MT, Ruhe J, Kroll S, Morhenn C, Kim S-T, Weigel D, et al. Microbial Hub Taxa Link Host and Abiotic Factors to Plant Microbiome Variation. PLoS Biol 2016; 14: e1002352.

55. Wang K, Senthil-Kumar M, Ryu C-M, Kang L, Mysore KS. Phytosterols play a key role in plant innate immunity against bacterial pathogens by regulating nutrient efflux into the apoplast. Plant Physiol 2012; 158: 1789–1802.

56. Moyes AB, Kueppers LM, Pett-Ridge J, Carper DL, Vandehey N, O’Neil J, et al. Evidence for foliar endophytic nitrogen fixation in a widely distributed subalpine conifer. New Phytol 2016; 210: 657–668.

57. Streubel J, Pesce C, Hutin M, Koebnik R. Five phylogenetically close rice SWEET genes confer TAL effector-mediated susceptibility to Xanthomonas oryzae pv. oryzae. New 2013.

58. Mayak S, Tirosh T, Glick BR. Plant growth-promoting bacteria that confer resistance to water stress in tomatoes and peppers. Plant Sci 2004; 166: 525–530.

59. Traxler MF, Kolter R. Natural products in soil microbe interactions and evolution. Nat Prod Rep 2015; 32: 956–970.

60. Barnard AML, Bowden SD, Burr T, Coulthurst SJ, Monson RE, Salmond GPC. Quorum sensing, virulence and secondary metabolite production in plant soft-rotting bacteria. Philos Trans R Soc Lond B Biol Sci 2007; 362: 1165–1183.

61. Stringlis IA, Zhang H, Pieterse CMJ, Bolton MD, de Jonge R. Microbial small molecules - weapons of plant subversion. Nat Prod Rep 2018; 35: 410–433.

62. Chelius MK, Triplett EW. The Diversity of Archaea and Bacteria in Association with the Roots of Zea mays L. Microb Ecol 2001; 41: 252–263.

63. Hodkinson BP, Lutzoni F. A microbiotic survey of lichen-associated bacteria reveals a new lineage from the Rhizobiales. Symbiosis 2009; 49: 163–180.

64. Huala E, Dickerman AW, Garcia-Hernandez M, Weems D, Reiser L, LaFond F, et al. The Arabidopsis Information Resource (TAIR): a comprehensive database and web-based information retrieval, analysis, and visualization system for a model plant. Nucleic Acids Res 2001; 29: 102–105.

65. Li H, Handsaker B, Wysoker A, Fennell T, Ruan J, Homer N, et al. The Sequence Alignment/Map format and SAMtools. Bioinformatics 2009; 25: 2078–2079.

66. Huson DH, Auch AF, Qi J, Schuster SC. MEGAN analysis of metagenomic data. Genome Res 2007; 17: 377–386.

67. diCenzo GC, Finan TM. The Divided Bacterial Genome: Structure, Function, and Evolution. Microbiol Mol Biol Rev 2017; 81: 00019–00017.

68. Mohanta TK, Bae H. The diversity of fungal genome. Biol Proced Online 2015; 17: 8.

69. Judelson HS. Dynamics and innovations within oomycete genomes: insights into biology, pathology, and evolution. Eukaryot Cell 2012; 11: 1304–1312.

70. Love MI, Huber W, Anders S. Moderated estimation of fold change and dispersion for RNA-seq data with DESeq2. Genome Biol 2014; 15: 550.

71. Friedman J, Alm EJ. Inferring correlation networks from genomic survey data. PLoS Comput Biol 2012; 8: e1002687.

72. Oksanen J, Kindt R, Legendre P, O’Hara B, Stevens MHH, Oksanen MJ, et al. The vegan package. Community ecology package 2007; 10: 631–637.

73. Choi K-H, Kumar A, Schweizer HP. A 10-min method for preparation of highly electrocompetent Pseudomonas aeruginosa cells: application for DNA fragment transfer between chromosomes and plasmid transformation. J Microbiol Methods 2006; 64: 391–397.

74. Exposito-Alonso M, Becker C, Schuenemann VJ, Reiter E, Setzer C, Slovak R, et al. The rate and potential relevance of new mutations in a colonizing plant lineage. PLoS Genet 2018; 14: e1007155.

75. Parker JE, Holub EB, Frost LN, Falk A, Gunn ND, Daniels MJ. Characterization of eds1, a mutation in Arabidopsis suppressing resistance to Peronospora parasitica specified by several different RPP genes. Plant Cell 1996; 8: 2033–2046.

